# Diffusion MRI Harmonization via Personalized Template Mapping

**DOI:** 10.1101/2023.05.12.540537

**Authors:** Yihao Xia, Yonggang Shi

## Abstract

One fundamental challenge in diffusion MRI (dMRI) harmonization is to disentangle the contributions of scanner-related effects from the variable brain anatomy for the observed imaging signals. Conventional harmonization methods rely on establishing an atlas space to resolve anatomical variability and generate a unified inter-site mapping function. However, this approach is limited in accounting for the misalignment of neuroanatomy that still widely persists even after registration, especially in regions close to cortical boundaries. To overcome this challenge, we propose a personalized framework in this paper to more effectively address the confounding from the misalignment of neuroanatomy in dMRI harmonization. Instead of using a common template representing site-effects for all subjects, the main novelty of our method is the adaptive computation of personalized templates for both source and target scanning sites to estimate the inter-site mapping function. We integrate our method with the rotation invariant spherical harmonics (RISH) features to achieve the harmonization of dMRI signals between sites. In our experiments, the proposed approach is applied to harmonize the dMRI data acquired from two scanning platforms: Siemens Prisma and GE MR750 from the Adolescent Brain Cognitive Development (ABCD) dataset and compared with a state-of-the-art method based on RISH features. Our results indicate that the proposed harmonization framework achieves superior performance not only in reducing inter-site variations due to scanner differences but also in preserving sex-related biological variability in original cohorts. Moreover, we assess the impact of harmonization on the estimation of fiber orientation distributions (FOD) and show the robustness of the personalized harmonization procedure in preserving the fiber orientation of original dMRI signals.

## 1. Introduction

Diffusion magnetic resonance imaging (dMRI) (Basser et al. 1994) allows the probing of brain microstructure *in vivo* and plays a key role in brain mapping research. With the widespread use of large-scale dMRI data from various studies (Mueller et al. 2005, Casey et al. 2018, Van Essen et al. 2013), the harmonization of dMRI data across acquisition protocols, sites, and vendors is a critical yet challenging problem (Vollmar et al. 2010, Zhu et al. 2011, Magnotta et al. 2012). One fundamental difficulty in dMRI harmonization is disentangling scanner-related effects from anatomical variations on the local appearance of imaging signals. To this end, we propose a novel personalized framework that better resolves the impact of anatomical variability on the estimation of inter-site mapping and advances the state-of-the-art in dMRI harmonization.

With the goal of limiting the impact of anatomical variability on the estimation of scanner effects, prevailing dMRI harmonization approaches rely on the co-registration of dMRI data across sites into a common atlas space. Then, spatial/regional mapping for dMRI harmonization can be estimated as the confounding of anatomy is expected to be removed. Following this registration-based framework, several statistical normalization approaches including Removal of Artificial Voxel Effect by Linear regression (RAVEL) (Fortin et al. 2016), Surrogate Variable Analysis (SVA) (Leek & Storey 2007), and ComBat (Johnson et al. 2007) were examined in (Fortin et al. 2017) for the harmonization of diffusion tensor imaging (DTI) features, and the ComBat method has been shown to be highly effective for multi-site DTI data pooling. For the direct harmonization of dMRI signals, the rotation invariant spherical harmonics (RISH) feature based method was proposed (Mirzaalian et al. 2015) and embedded in a registration framework to achieve voxel-wise harmonization in a template space (Mirzaalian et al. 2018, Cetin Karayumak et al. 2019). Similarly, a method of moments (MoM) was proposed (Huynh et al. 2019) to achieve the harmonization of dMRI signals after registration to an atlas space by comparing the spherical moments of dMRI signals across sites to estimate a linear mapping function.

Despite the essential role of co-registration in previous harmonization methods, little work has been done to examine its reliability in building anatomical correspondences across subjects in dMRI harmonization. In fact, the inter-subject variability in neuroanatomy and hence the possible lack of one-to-one correspondences across subjects will inevitably complicate the construction of anatomical correspondences essential for the validity of existing harmonization models. This difficulty will be especially evident in regions surrounding the cortical boundaries, where the high variability across subjects has been well-known (Thompson et al. 1996, Uylings et al. 2005). As illustrated in Figure 1(a), there are obvious anatomical differences between the two Adolescent Brain Cognitive Development (ABCD) (Casey et al. 2018) subjects even after they have been co-registered to the template image in Figure 1(b). Because harmonization algorithms rely on pooling data across subjects from different sites at locations with *corresponding* anatomy, this type of mismatch, shown in Figure 1, would make the estimation of the inter-site mapping function much less reliable and lead to invalid harmonization of the dMRI data at affected locations.

**Figure 1:**
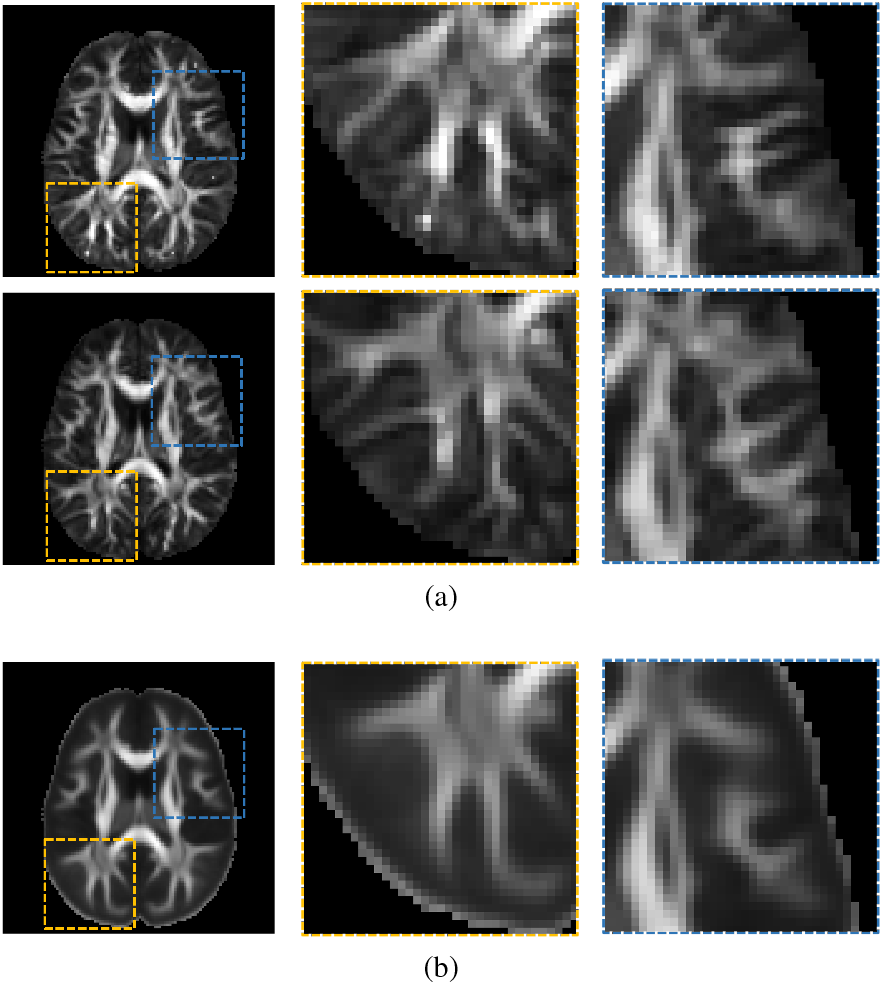
Inter-subject anatomical variability and the resulting mismatch between individual subjects and the sample mean template. (a) FA images of two ABCD subjects co-registered to the template space. Top: NDARINV15MFU6UZ acquired from a GE MR750 scanner; bottom: NDARINV1NW3HM13 scanned on a Siemens Prisma scanner. (b) A template image constructed from the FA images of 100 ABCD subjects. Left column: axial slices of the two ABCD subjects and the template. For each axial slice, zoomed views of the ROIs highlighted by yellow and blue boxes are shown in the middle and right columns, respectively, where the variations of the brain anatomy are noticeable.

To overcome this fundamental problem in establishing anatomically consistent correspondences for the estimation of inter-site mapping functions, we develop a personalized template estimation framework in this paper to abridge the anatomical gaps between population-based templates and individual subjects to be harmonized. We also integrate this personalized analysis framework with RISH features (Mirzaalian et al. 2018, Cetin Karayumak et al. 2019) to achieve voxel-wise harmonization of dMRI signals across scanners. Figure 2(a) shows the over-all harmonization framework. To integrate the feature representations of reference subjects from both source and target scanning sites for each query subject, we introduce a personalized pooling tensor for each site. As illustrated in Figure 2(b), the pooling tensor of a site consists of weight vectors computed via solving a convex optimization problem at each voxel based on the similarity of the local anatomy between the query subject and the reference subjects, which are representative of a site. Once the pooling tensors are estimated, site-specific templates will be estimated in a personalized manner for computing the inter-site mapping function for each feature representation. By combining the transferred feature representations, we achieve personalized harmonization of the dMRI data from the source to the target site for the query subject.

**Figure 2:**
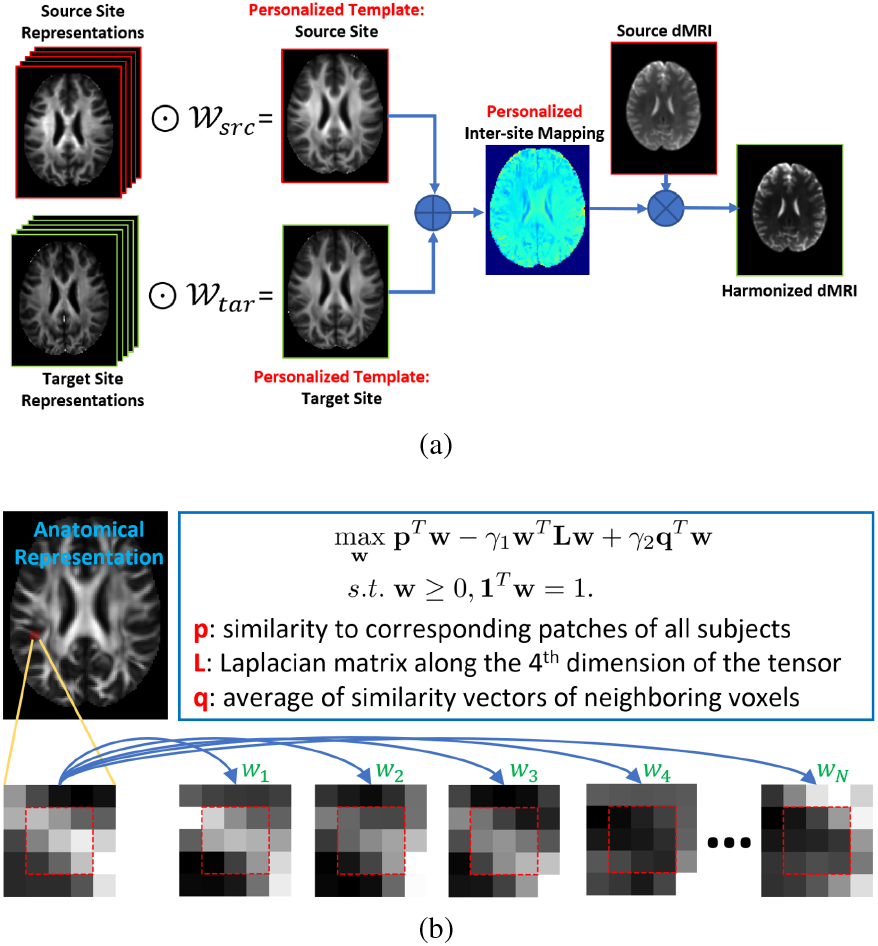
An overview of our personalized dMRI harmonization method. (a) The overall framework for the estimation of personalized templates and inter-site mapping functions to achieve the harmonization from the source to the target site. (b) Details about the estimation of the personalized pooling tensors (**𝒲** _*src*_ and **𝒲** _*tar*_) by computing a weight vector *w* at each voxel as the solution of a convex optimization problem based on local similarity of the anatomy between the query subject and reference subjects of each site.

The rest of this paper is organized as follows. Section 2 introduces the personalized dMRI harmonization framework and related numerical algorithms. Experimental results are presented in Section 3 to demonstrate that the proposed personalized method is able to achieve much-improved performance in comparison with the state-of-the-art method based on RISH features mapping. Finally, discussions and conclusions are made in Section 4.

## 2. Methods

### 2.1. Diffusion MRI harmonization in a common space

Given dMRI data from a source and target site, the goal of the harmonization task is to estimate a mapping function: *Ŝ* = Ψ(*S*) for each subject in the source site, where *S* denotes the diffusion signal of a subject from the source site and Ψ denotes the inter-site mapping function to the target site. For the estimation of the inter-site mapping function, a set of reference subjects from the source and target sites are typically needed to represent the scanner-dependent variations. In addition, it is important to consider the anatomical variations across subjects and the spatial heterogeneity of scanner differences.

To this end, nonlinear image registration has played a key role in minimizing the impact of anatomical differences across subjects and allowing the pooling of information from representative reference subjects for the characterization of site-effects. Within a common space constructed through nonlinear registration of the anatomical images of subjects from both sites, the harmonization mapping function Ψ can be determined by comparing the rotation invariant diffusion features, such as RISH features (Cetin Karayumak et al. 2019) and diffusion moments (Huynh et al. 2019), across sites. For each feature, a population-based template **E** is first constructed for both the source and target sites by evenly pooling the co-registered feature images of the reference subjects from each site:

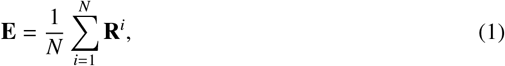

where **R**^*i*^ is the diffusion feature image of the i-th reference subject from a given site. Using the population-based feature templates from both the source and target site, the harmonization mapping function Ψ can be estimated at each location in the common space, which can then be applied to all source subjects for their harmonization to the target site (Cetin Karayumak et al. 2019, Huynh et al. 2019).

### 2.2. Personalized template estimation

While the common space from nonlinear registration can greatly reduce inter-subject variations, significant anatomical differences remain, especially around the cortical areas. For a given query subject from the source site to be harmonized, this type of anatomical misalignment will no doubt introduce significant bias to the estimation of feature templates and consequently affect the harmonization mapping function. To avoid the confounding caused by the anatomical misalignment in harmonization, we develop a personalized approach for template estimation by adaptively integrating the feature representations from the reference subjects of a given site in the common space according to localized anatomical similarity. For a given query subject **𝒬** from the source site, we compute a personalized pooling tensor **𝒲**^**𝒬**^ for the estimation of a personalized template **E**^**𝒬**^:

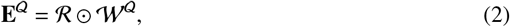

where **ℛ** = [**R**^1^ … **R**^*N*^] denotes the feature images of *N* reference subjects from a given site. At a location *x* in the common space, the uneven pooling of references is performed by a weighted sum:

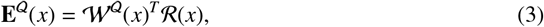

where **𝒲**^**𝒬**^ (*x*) = [*w*_1_(*x*), …, *w*_*N*_ (*x*)]^*T*^ is a weight vector, with *w*_*i*_(*x*) > 0 for *i* = 1, …, *N* and 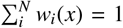 and **ℛ** (*x*) = [**R**^1^(*x*), …, **R**^*N*^ (*x*)]^*T*^ is the local reference vector containing diffusion features sampled from the reference subjects.

For the calculation of the weight tensor **𝒲**^**𝒬**^, we compute the reference weight vector **𝒲**^**𝒬**^(*x*) adaptively at each location *x* in the common space by solving a convex optimization problem. Let’s denote that the co-registered anatomical image of the source query subject in the common *σ*_*U*_ *σ*_*V*_ space as *U* and those of the *N* reference subjects as **𝒱** = {*V*_1_, *V*_2_, …, *V*_*N*_ }. For numerical implementation, we use fractional anisotropy (FA) image as the anatomical image of each subject in this work. At each point *x* in the common space, the local anatomical similarity measure between the query subject and a reference subject is denoted as *sim*(*U*(*x*), *V*(*x*)), which we define based on the cross-correlation of local patches: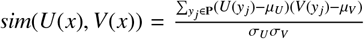, where **P** is a 3D patch of size 3 × 3 × 3 centered at the voxel *x* in both *U* and *V, μ*_*U*_ and *μ*_*V*_ are the mean FA value within the patch, and *σ*_*U*_ and *σ*_*V*_ are the standard deviations. To ensure that the personalized templates are representative of reference subjects with similar anatomy at each point, we introduce a Laplacian regularizer across reference subjects in the cost function. This regularizer can encourage the reference subjects with highly similar anatomy to be assigned with similar weights. Furthermore, we include a spatial consistency term in the cost function to consider the local spatial dependencies in adjacent locations. The weight vector estimation is then formulated as the following convex optimization problem:

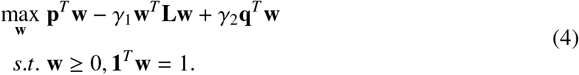

where **w** = **𝒲** ^**𝒬**^ (*x*) is the weight vector at *x* and **p** = [*sim*(*U*(*x*), *V*_1_(*x*)), …, *sim*(*U*(*x*), *V*_*N*_ (*x*))]^*T*^ is the query-to-reference similarity vector at *x*. In the Laplacian regularization term **w**^*T*^ **Lw, L** = **D** − **A** is a Laplacian matrix, where **A** is the inter-reference similarity matrix with entry **A**(*i, j*) = *sim*(*V*_*i*_(*x*), *V*_*j*_(*x*)), and **D** is the degree matrix. We can trade off between the potential mismatch of brain anatomy and the reliability in template estimation by tuning the parameter *γ*_1_. By increasing the parameter *γ*_1_, the weights would be propagated to more reference subjects, which would enlarge the sample size and improve the reliability of the personalized templates. On the other hand, an overly large parameter *γ*_1_ could result in assigning non-negligible weights to all reference subjects and bring the misregistration problem back to template construction. The third term of the cost function is designed to encourage the spatial consistency of the weight vectors, where the neighborhood similarity vector for the query location *x* is defined as

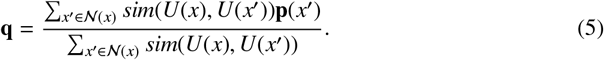

Here **𝒩** (*x*) is the neighborhood centered at *x* and **p**(*x*^′^) = [*sim*(*U*(*x*^′^), *V*_1_(*x*^′^)), …, *sim*(*U*(*x*^′^), *V*_*N*_ (*x*^′^))]^*T*^ is the query-to-reference similarity vector of the neighboring voxel *x*^*′*^ ∈ **𝒩** (*x*). Note that the contributions of spatial neighborhoods are proportional to their anatomical similarity with the query location to accommodate the anatomical inconsistencies that may arise in neighborhood patches located at tissue boundaries. The tuning parameter *γ*_2_ controls the involvement of neighborhood information in the estimation of weight assignment. Overall, this quadratic programming problem is solved by using the ECOS solver (Domahidi et al. 2013, Diamond & Boyd 2016).

For dMRI harmonization, personalized templates will be computed for each query subject with respect to the reference subjects from both the source and target site, respectively. This procedure will be performed for all pertinent diffusion image features which characterize the scanner effects for the source and target sites. Subsequently, the inter-site mapping function for harmonizing the dMRI signal to the target site can be estimated, as we will describe below.

### 2.3 Personalized harmonization based on mapping spherical harmonics features

To demonstrate the application of our personalized template estimation for dMRI harmonization, we integrate it with the rotational invariant spherical harmonics (RISH) features from the LinearRISH harmonization method (Cetin Karayumak et al. 2019). To compute the RISH features, the dMRI signals at each voxel are first represented by the spherical harmonics (SPHARM) as *S* ≃ ∑_*lm*_ *C*_*lm*_*Y*_*lm*_, where *Y*_*lm*_ and *C*_*lm*_ are the SPHARM basis and corresponding coefficient of order *l* and degree *m*. The RISH features characterize the energy distribution of the dMRI signal at each order *l* and are defined as 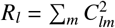. In a common space constructed by nonlinear registration, the LinearRISH method computes a template from the source and target site for each RISH feature and calculates a scale factor **Φ**_*l*_ at each location, which is then applied to scale the SPHARM coefficients at the same order *l*: *Ĉ*_*lm*_ = Φ_*l*_*C*_*lm*_ to realize the harmonization. Note that the template and scale factors are the same for all subjects from the source site.

For the personalized harmonization of dMRI data from a source site (*src*) to a target site (*tar*) in our proposed method, the mapping function **Φ**_*l*_ is estimated separately for each query subject **𝒬** at each SPHARM order:

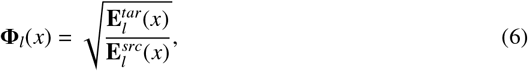

where 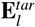 and 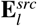 are the personalized template of the RISH feature at the order *l* for the target *l* and source sites, respectively. Compared with the conventional LinearRISH method, the main novelty of our method is that the RISH feature templates are estimated by solving (4) and hence emphasize reference subjects with similar anatomy to boost consistency. The mapping image **Φ**_*l*_ is then warped back to the subject space of the query individual. At each voxel in the query space, the SPHARM coefficient 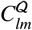 of the dMRI signal is scaled by the mapping function at the order *l* as 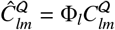. Finally, the harmonized dMRI signal *Ŝ* ^𝒬^ is generated by using the scaled

### SPHARM coefficients

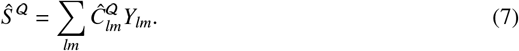

By repeating this process for all voxels, we obtain the harmonization of the query subject from the source to the target site.

## 3. Experimental Results

To demonstrate the efficacy of the proposed personalized dMRI harmonization framework, we apply the proposed method to harmonize dMRI data acquired from two scanning systems: Siemens Prisma and GE MR750 from the ABCD study (Casey et al. 2018) and compare the performance with the state-of-the-art LinearRISH method (Cetin Karayumak et al. 2019). The results are organized as follows. In Section 3.1, we describe the dMRI dataset and the implementation details of harmonization experiments. An illustration of the weight assignment for personalized template construction in our method and comparison with the template from LinearRISH are presented in Section 3.2. Next, we discuss the selection of regularization parameters for our method in Section 3.3. In Section 3.4, we present a comprehensive comparison of the proposed harmonization framework with LinearRISH in terms of reducing the inter-site variation of DTI features. Then, we assess the impact of available reference subjects on the performance of our personalized harmonization framework in Section 3.5. In Section 3.6, we focus on the regional distribution of DTI features in cortical gray matter areas to further demonstrate the effectiveness of our personalized harmonization method in disentangling site-effects from variable brain anatomy. In Section 3.7, we examine the impact of dMRI harmonization on the estimation of fiber orientation distribution. Finally, we investigate the preservation of sex-related biological variability after harmonization and compare the performance of both methods in Section 3.8.

### 3.1. Dataset and implementation details

#### 3.1.1. Subjects and dMRI data

In our experiments, we used data acquired on two scanning systems: Siemens Prisma (SIEMENS) and GE Discovery MR750 (GE) from the ABCD Study (Casey et al. 2018). A total of 200 subjects at baseline were selected from the ABCD study, with balanced sample sizes for each scanner, including 100 subjects scanned on Siemens Prisma and another 100 subjects on GE Discovery MR750, which we denote as the SIEMENS and GE cohort in our experiments, respectively. To control for confounding effects of age and gender, all selected participants are 9 years old and well-matched for gender, consisting of 50 females and 50 males in each cohort. For both scanners, the ABCD dMRI data used in harmonization experiments were acquired at b = 3000 *s*/*mm*^2^ from 60 gradient directions using multiband EPI (Kawin Setsompop 1 et al. 2011). All dMRI data have an isotropic spatial resolution of 1.7 *mm*. The sequence parameters, repetition time (TR), time to echo (TE), and flip angle, are scanner-specific. See Table 1 for detailed dMRI sequence parameters for the SIEMENS and GE scanners.

**Table 1:** Diffusion MRI parameters for the Siemens Prisma and GE 750 scanner in the ABCD study.

| Scanner | TR (ms) | TE (ms) | Flip Angle (°) |
| --- | --- | --- | --- |
| Siemens | 4100 | 88 | 90 |
| GE | 4100 | 81.9 | 77 |

Furthermore, all data used in our experiments are part of the ABCD Data Release 4.0 from the NIMH Data Archive (NDA) and have been preprocessed by the ABCD study. Quality control and preprocessing were performed to minimize the influence of poor image quality and artifacts prior to harmonization (Hagler Jr et al. 2019). For eddy current correction, distortions on the diffusion gradients were estimated using the approach in (Zhuang et al. 2006). The echo-planar imaging distortions were corrected for each acquisition using the reversing gradient method with FSL’s TOPUP (Andersson et al. 2003; 2012). The head motion correction and diffusion gradient adjustment were performed following procedures in (Hagler Jr et al. 2009). Additionally, to minimize the impact of noise on the RISH based harmonization (Cetin Karayumak et al. 2019, Mirzaalian et al. 2016), the diffusion-weighted images were denoised via the over-complete local PCA-based method (Manjón et al. 2013).

#### 3.1.2. Implementation details of dMRI harmonization

For both our method and the LinearRISH method for comparison, we calculated five RISH features for SPHARM orders of *l* = {0, 2, 4, 6, 8} for the harmonization of dMRI signals at each voxel. For the generation of the common space for computing the harmonization mapping function, we applied the template construction tool from Advanced Normalization Tools (ANTs) (Avants et al. 2009) to the FA images of all subjects. The generated deformation fields were used to perform inter-subject alignment of the RISH and DTI features in all harmonization and evaluation experiments. For our method, all subjects from the source or target site (SIEMENS or GE) served as reference subjects for the construction of personalized RISH feature template and the estimation of mapping functions.

### 3.2. Local reference selection and personalized template construction

Personalized template construction is achieved by adaptively assigning pooling weights at each voxel according to the anatomical similarities between the query subject and reference subjects from each site. First, we illustrate how the weight assignment was conducted at two typical locations with parameters: *γ*_1_ = 0.1 and *γ*_2_ = 2.0. Figure 3(a) presents a location centered in a white matter region where the anatomical mismatching problem is relatively mild. In this example, both the query subject and reference subjects were from the GE cohort. As can be seen in the top row on the right of Figure 3(a), reference patches with high anatomical similarity were assigned high weights and included for template construction. On the other hand, reference patches with low similarity (shown in the bottom row of Figure 3(a)) were assigned zero weights and excluded from template estimation at the current voxel. Figure 3(b) shows another example at the boundary of gray and white matter with higher inter-subject anatomical variability. Here, the query subject was from the GE cohort and the reference subjects were from the SIEMENS cohort. The top row on the right of Figure 3(b) presents the instances of selected reference patches with non-zero weights. Compared to the results shown in Figure 3(a), it is observable that the pooling weights are more concentrated over well-aligned references because a more limited number of references share similar anatomy to the query subject at the highlighted location on the cortical boundary. As shown in the bottom row on the right of Figure 3(b), noticeable dissimilarities exist between the query and reference patches. With zero weights assigned to those references, our method effectively alleviates the confounding due to the misalignment of brain anatomy in template construction.

**Figure 3:**
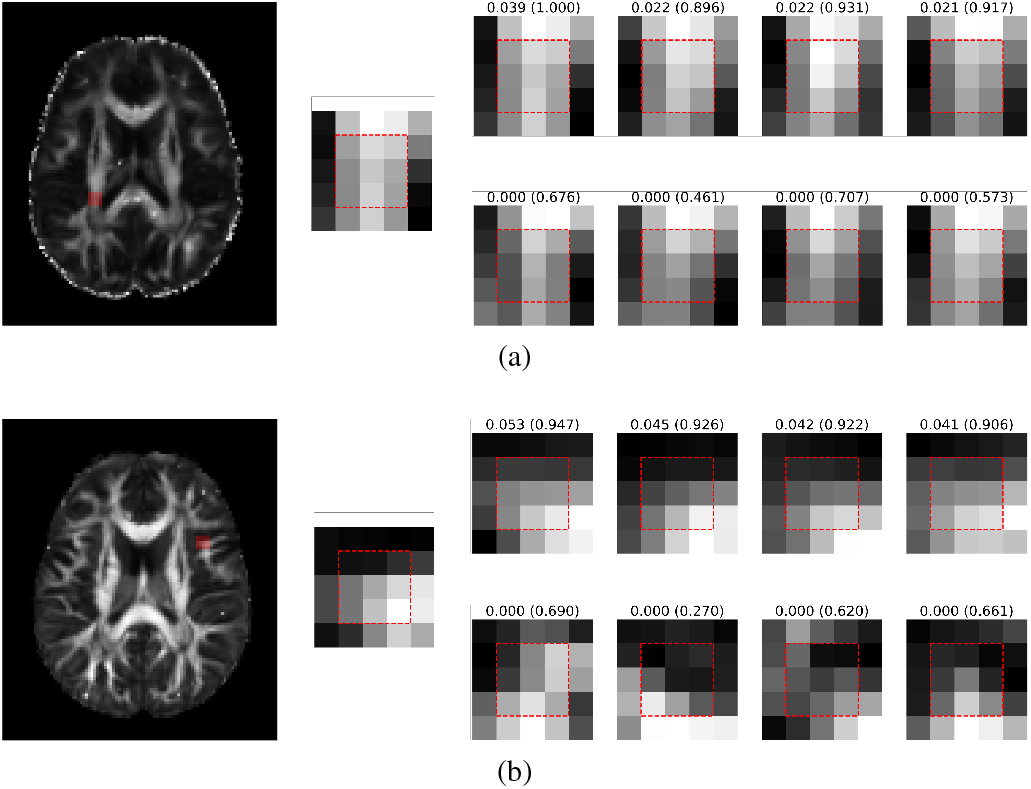
An illustration of selected and discarded references at two representative locations. In each subfigure (a) and (b), the results are organized as follows. Left: the transverse slice of the FA image indicating the location of the query patch in the common space (highlighted in red); middle: a zoomed view of the query patch (in red box); right (top row): top-4 reference patches ordered by their pooling weights; right (bottom row): discarded reference patches with zero weights. The assigned weights and the query-reference anatomical similarity are displayed above each reference patch in the form of “weight (anatomical similarity)”.

By performing the personalized weight assignment at each voxel for reference subjects from the source or target site, personalized templates can be generated for a given query subject to characterize site effects in dMRI signals. Figure 4 displays the zeroth-order RISH feature of a query subject (Subject ID: NDARINV15MFU6UZ) and the corresponding templates estimated by the LinearRISH method based on sample mean and the proposed personalized method. In contrast to the conventional templates from LinearRISH (first row on the right) for both the GE and SIEMENS cohort, we can see the personalized templates from our method (second row on the right) capture much more cortical folding details (circled by red dashed lines) that align very well with the query subject.

**Figure 4:**
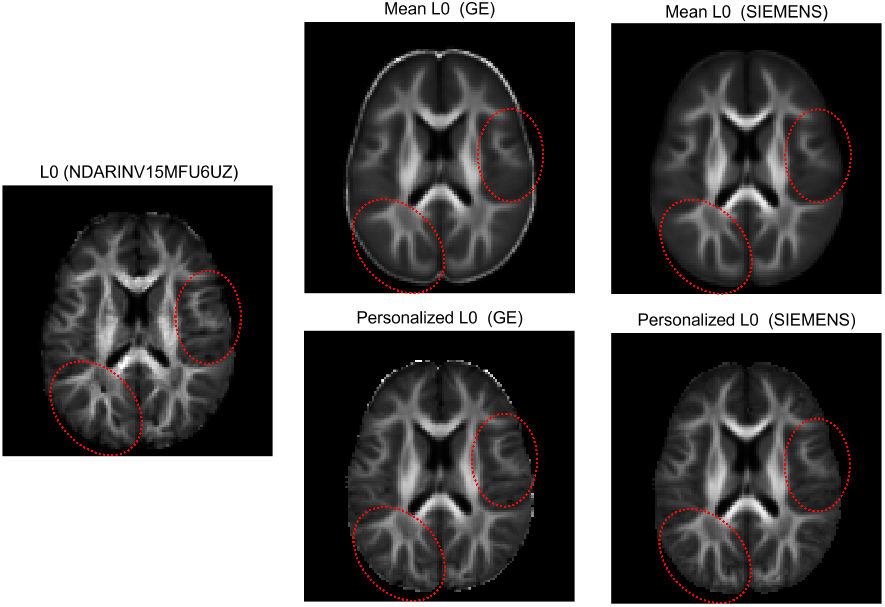
A comparison of site-specific templates estimated by the LinearRISH method based on sample mean and our personalized method for the zeroth-order RISH feature. Left: a transverse slice of the zeroth-order RISH feature of a query subject (GE Subject ID: NDARINV15MFU6UZ). Right (top row): the corresponding slice of templates estimated from the GE and SIEMENS cohort by the LinearRISH method. Right (bottom row): the corresponding slice of the templates estimated from the GE and SIEMENS cohort by our personalized method.

### 3.3. Regularization parameters

#### 3.3.1. Across-reference regularization

In the proposed personalized harmonization framework, the number of selected references primarily depends on the underlying query-to-reference anatomical similarity at each voxel and the choice of *γ*_1_. We investigated the spatial distribution of the average number of selected references per query subject with varying *γ*_1_ while keeping *γ*_2_ = 2.0. Because anatomical similarity across subjects varies in different brain regions, the average number of references used for personalized template estimation differs spatially. For two harmonization experiments: GE toward SIEMENS and SIEMENS toward GE, the spatial distribution of reference number for the estimation of site-specific templates are plotted in Figure 5 (a) and (b) under different parameter choices of *γ*_1_. From the results, we can observe that white matter areas consistently have a higher reference number than cortical gray matter regions. This suggests that gray matter regions with higher inter-subject variability tend to have fewer qualified references. Furthermore, an increase of *γ*_1_ enlarges the size of the reference set. By tuning *γ*_1_, we trade off between the potential mismatch to references and the reliability achieved through the availability of a sufficient number of references to represent site-specific characteristics at each location. Upon examining the distribution of reference numbers with *γ*_1_ = 0.1, the number of references typically exceeds 20 in white matter regions, which is more than sufficient for a reliable representation of population characteristics in harmonization (Cetin Karayumak et al. 2019). For gray matter regions with more severe misalignment in brain anatomy, there are still on average at least 10-15 references available for template construction. Based on these observations, we set *γ*_1_ = 0.1 for the rest of our experiments.

**Figure 5:**
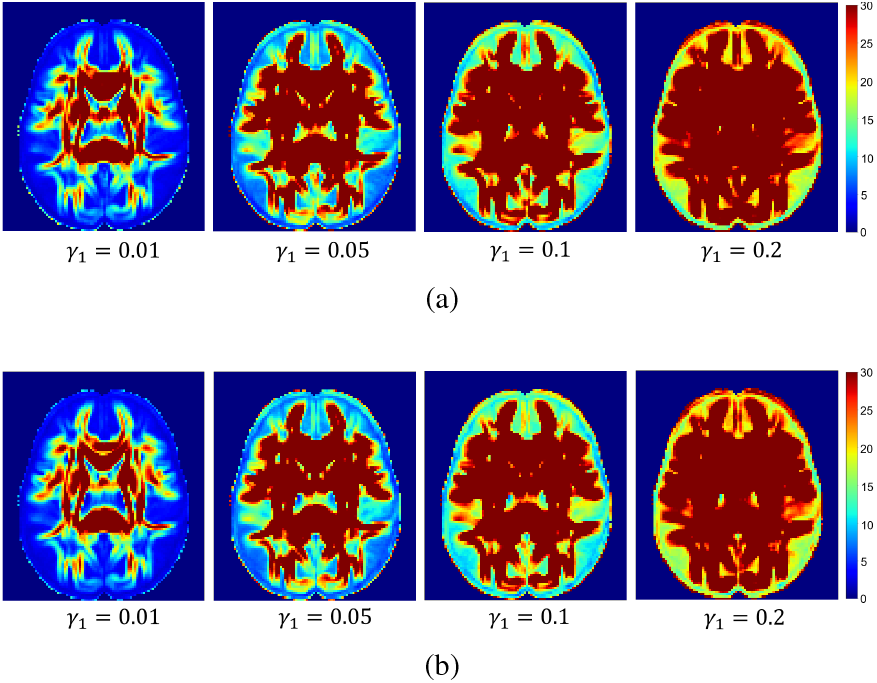
The spatial distribution of reference numbers in personalized and site-specific template construction with different choices of the regularization parameter *γ*_1_. (a) Query subjects from the GE cohort and Reference subjects from the SIEMENS cohort. (b) Query subjects from the SIEMENS cohort and Reference subjects from the GE cohort.

#### 3.3.2 Spatial smoothness regularization

We further examined the effect of the regularization parameter *γ*_2_ on the spatial consistency of reference selection. For each query subject **𝒬**, the 4D personalized pooling tensor **𝒲**^**𝒬**^is composed of multiple 3D weight images 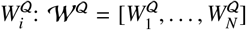, where *N* is the size of the reference set from a site. For each weight image, we used the magnitude of its Laplacian to quantify the smoothness of weight assignment (Xue et al. 2014). Overall, the spatial smoothness of reference selection for a query subject is computed as follows:

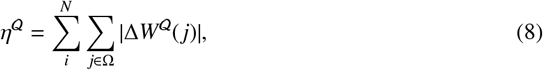

where 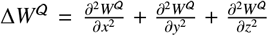 is the Laplacian of a 3D weight image and Ω denotes the location set of the whole brain region in the common space. In Figure 6, we demonstrate the relationship between the parameter *γ*_2_ and the overall smoothness of weight assignment in situations where query and reference subjects are from different sites while maintaining *γ*_1_ = 0.1. It can be observed that increasing *γ*_2_ leads to a noticeable improvement in the spatial smoothness of weight assignment. After *γ*_2_ reaches 1.0, its impact on the smoothness reaches a region of relative stability. The turning point is at *γ*_2_ = 2.0 and larger values of *γ*_2_ result in a tendency towards less spatial consistency. Thus, it is reasonable to choose *γ*_2_ around 2.0. In the following experiments, we set *γ*_2_ = 2.0 for the regularization of spatial consistency.

**Figure 6:**
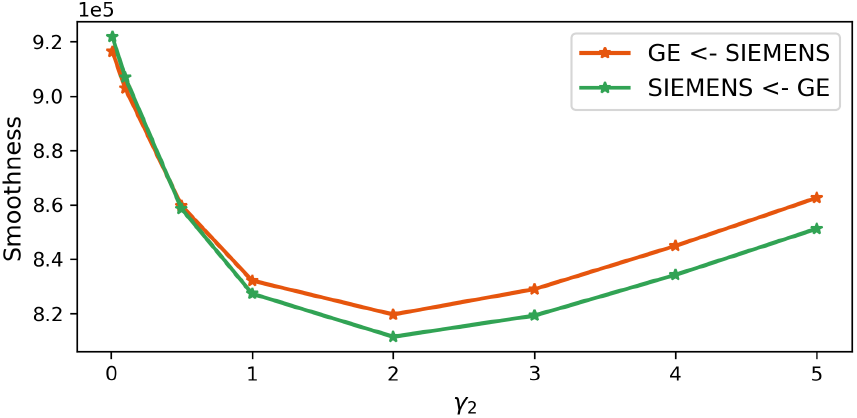
Spatial consistency regularization by changing the parameter *γ*_2_. For query subjects from the GE cohort and reference subjects from the SIEMENS cohort, the mean values of the weight smoothness measure across all query subjects with respect to varying *γ*_2_ are plotted in red. Similarly, the green curve shows the results under an alternative setting (query subjects from the SIEMENS and reference set from the GE cohort).

### 3.4. Inter-site variation and harmonization

To quantitatively evaluate the performance of harmonization methods, we calculated the inter-site coefficients of variation (CoVs) for diffusion features, including fractional anisotropy (FA) and mean diffusivity (MD), before and after harmonization. The coefficient of variation, defined as the ratio of the standard deviation to the mean of a feature, quantifies the dispersion of a distribution. The inter-site CoVs, reflecting the discrepancy of integrated datasets, are expected to be reduced after harmonization. For a qualitative evaluation, Figure 7 presents the voxel-wise CoVs of merged dMRI data for the following combinations: (1) original SIEMENS and GE data (SIEMENS + GE), (2) original SIEMENS and harmonized GE data using SIEMENS as the target site (SIEMENS + harmonized GE), and (3) original GE and harmonized SIEMENS data using GE as the target site (GE + harmonized SIEMENS).

**Figure 7:**
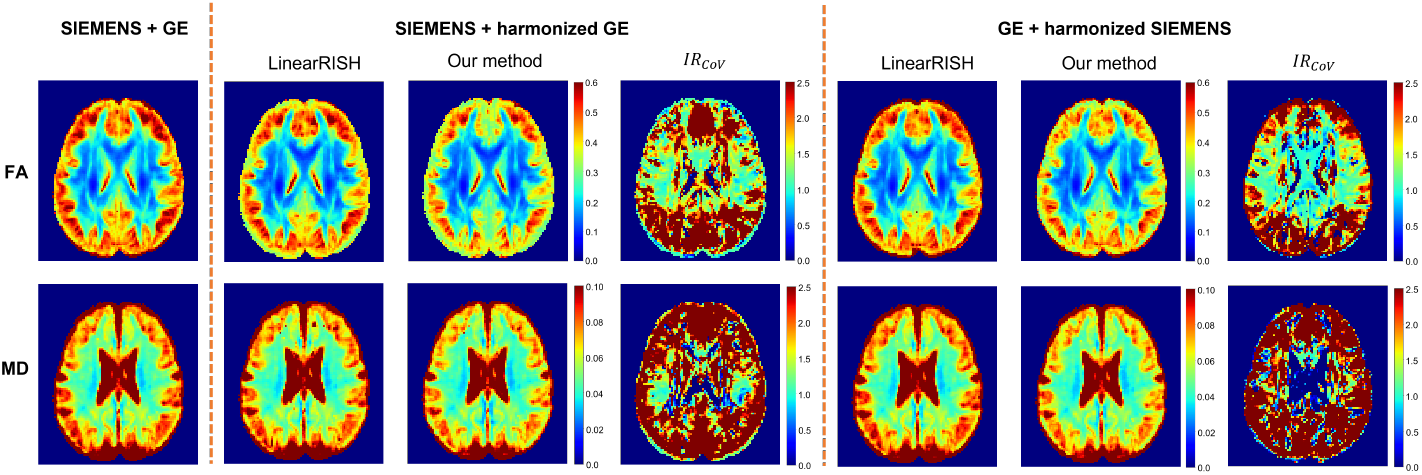
Inter-site coefficients of variation of FA and MD before and after harmonization. The results for original datasets before harmonization, SIEMENS + harmonized GE, and GE + harmonized SIEMENS are shown in the left, middle, and right panel. In the second and third panel, the results of LinearRISH, our method, and the improvement rate (*IR*_*CoV*_) are displayed from left to right, respectively.

To clearly demonstrate the improvement achieved by our proposed method, we calculated the local improvement rate (*IR*_*CoV*_) defined as follows:

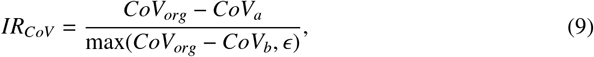

where *CoV*_*org*_, *CoV*_*b*_, and *CoV*_*a*_ were the CoVs before harmonization, after harmonization using the baseline method (LinearRISH), and using the proposed method. To ensure a positive denominator, ϵ was set to be 0.001. A value of *IR*_*CoV*_ > 1 indicates superior performance by our method in reducing the CoV of the integrated data from the GE and SIEMENS cohort. The voxel-wise *IR*_*CoV*_ images for FA and MD are shown in the fourth and last column in Figure 7 for two harmonization tasks: harmonizing GE data to SIEMENS (SIEMENS + harmonized GE) and harmonizing SIEMENS data to GE (GE + harmonized SIEMENS). We can clearly see that our method achieved better performance than LinearRISH in most brain regions, especially in areas close to cortical gray matter. In other white matter regions, the two methods achieve comparable performance in both harmonization tasks.

Besides, we calculated the average CoV across the whole brain and the negative rate (*r*_*N*_) of CoV after harmonization to quantify the overall performance of the two harmonization methods. The negative rate (*r*_*N*_) of CoV is defined as the ratio of the number of voxels with increasing or equal CoV after harmonization to the total number of voxels in the common space. It represents the failure rate in terms of reducing the inter-site variability. Table 2 summarizes the quantitative results for both harmonization tasks: SIEMENS + harmonized GE and GE + harmonized SIEMENS using the proposed method and LinearRISH. This further demonstrates the much improved performance by our personalized harmonization framework in reducing the inter-site variation and the negative rate (*r*_*N*_) for both the FA and MD feature. and (d) show the corresponding results for GE + harmonized SIEMENS data. The inter-site CoVs and the negative rates of CoV for the results produced by LinearRISH with 100 references from each platform are in red dashed-line and blue dashed-line, respectively, in each panel.

**Table 2:** Inter-site coefficients of variation and negative rates (*r*_*N*_) after harmonization for FA and MD.

| Dataset | FA CoV ($r_N$) | MD CoV ($r_N$) |
| --- | --- | --- |
| SIEMENS+GE | $0.404 \pm 0.160$ (-) | $0.090 \pm 0.052$ (-) |
| SIEMENS+harmonized GE (LinearRISH) | $0.361 \pm 0.132$ (9.83%) | $0.087 \pm 0.047$ (15.47%) |
| SIEMENS+harmonized GE (our method) | $0.332 \pm 0.121$ (2.27%) | $0.080 \pm 0.043$ (3.14%) |
| GE+harmonized SIEMENS (LinearRISH) | $0.409 \pm 0.172$ (30.04%) | $0.096 \pm 0.064$ (65.71%) |
| GE+harmonized SIEMENS (our method) | $0.367 \pm 0.150$ (11.58%) | $0.088 \pm 0.055$ (15.56%) |

### 3.5. Number of available references

When constructing personalized templates in our method, having an adequate number of reference subjects from each site is beneficial to cover the variability of neuroanatomy, especially in regions where the anatomical structures are not shared by the majority of the population. In this experiment, we assessed the impact of the number of available references on harmonization performance. The same parameter settings were used with *γ*_1_ = 0.1 and *γ*_2_ = 2.0. For both the GE and SIEMENS scanners, we randomly selected subsets, varying in size from 20 to 100, of the GE and SIEMENS cohorts as references for personalized template construction. For both the GE and SIEMENS scanners, the same number of references were used in all experiments. Figure 8 shows that the inter-site CoVs (in red) and the negative rates (*r*_*N*_) of CoV (in blue) for both FA and MD decrease as the size of the reference set increases for both harmonization tasks. This indicates that sufficient references (≥ 60) are crucial for adaptive template construction and the success of dMRI harmonization.

**Figure 8:**
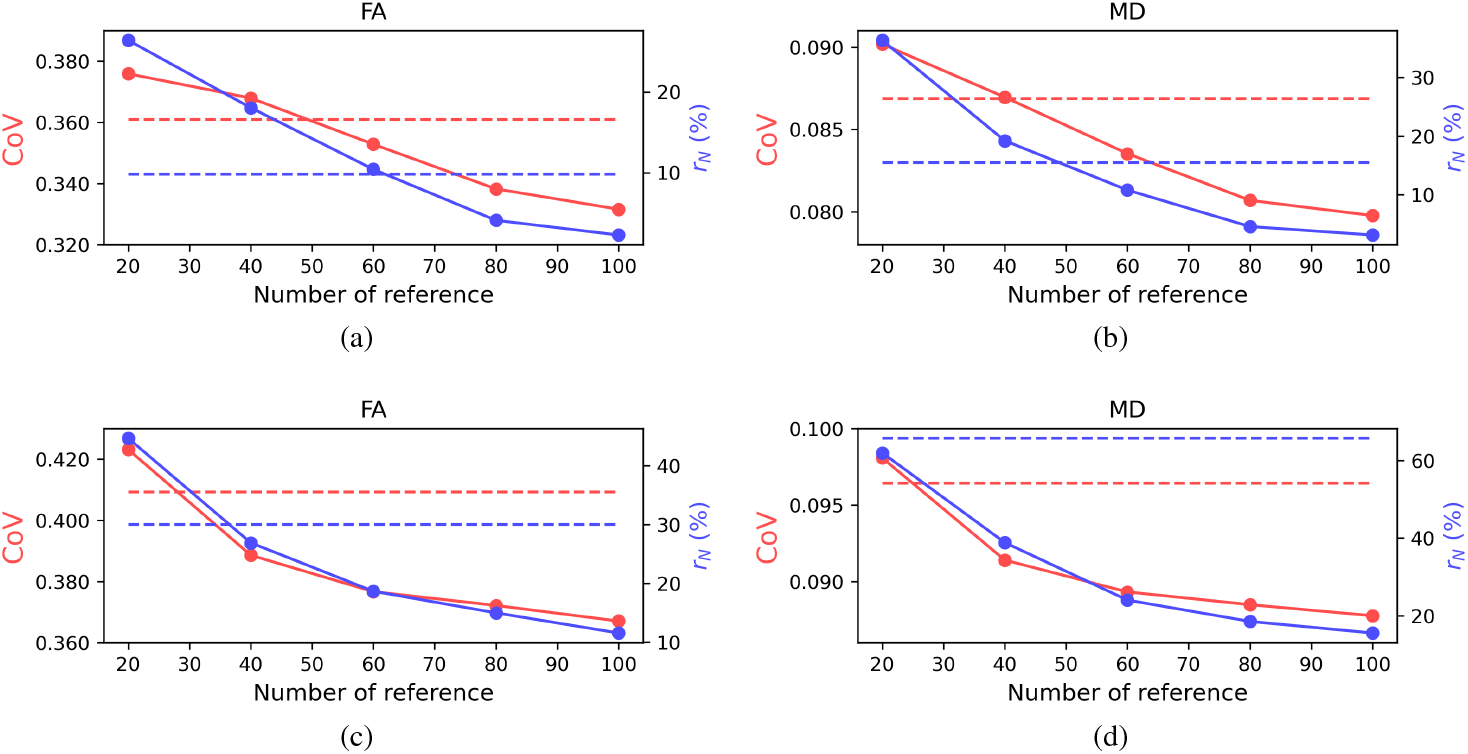
Inter-site CoVs for FA and MD with varying numbers of available references. (a) and (b) display the inter-site CoVs (in red line) and the negative rates of CoV (in blue line) of FA and MD with SIEMENS + harmonized GE data.

### 3.6. Impact of personalized harmonization on DTI feature distribution in cortical gray matter regions

In section 3.4, we presented the overall performance enhancement achieved by our method and qualitatively visualized the major improvement in cortical gray matter regions. To further assess the performance of harmonization, we analyzed the distribution of DTI features in cortical lobes before and after harmonization. As we noticed that the task of harmonizing GE toward SIEMENS data achieved lower inter-site CoVs in section 3.4, we focused on DTI features computed from harmonized GE data toward SIEMENS by our method and LinearRISH. As shown in Figure 9, four histograms of the FA feature (SIEMENS, GE, harmonized GE by our method, harmonized GE by LinearRISH) within the gray matter voxels of each cortical lobe (frontal, pariental, temporal, and occipital) of the left hemisphere are plotted. The cortical lobes were defined by merging parcellated ROIs from FreeSurfer (Fischl 2012). By comparing the respective FA feature distribution curves in Figure 9, we observe that both our method and LinearRISH can reduce the noticeable discrepancies between the original GE (blue curves) and SIEMENS (red curves) data before harmonization. However, it is evident that our method achieves better performance as the FA histograms of harmonized data using our method (purple curves) align more closely with the target histograms (red) than those produced by LinearRISH (cyan curves). We can quantify the difference of feature distributions across scanners using the Jensen–Shannon divergence (JSD) (Lin 1991):

**Figure 9:**
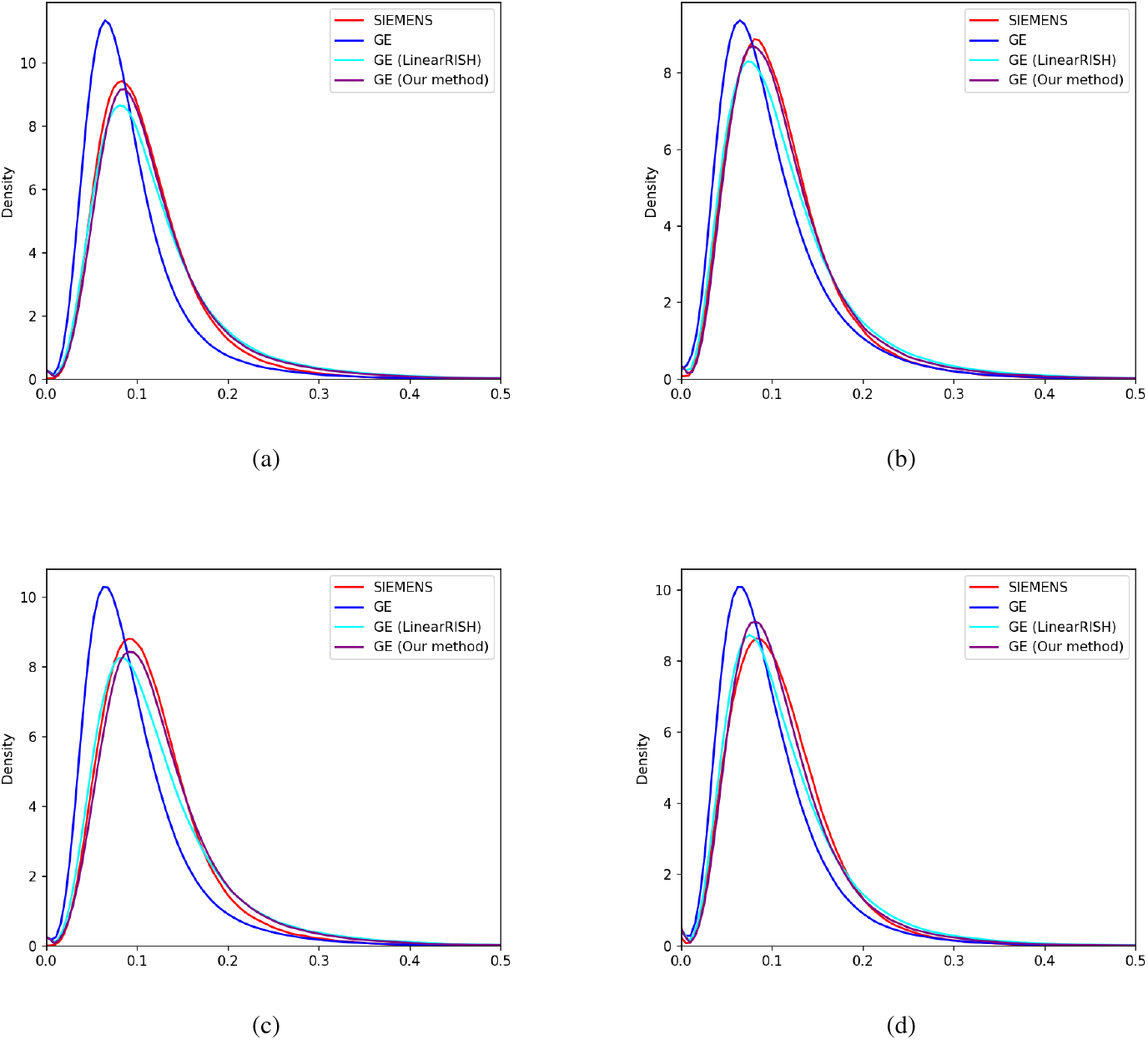
FA distributions before and after harmonization. The density curves of FA within the frontal, occipital, parietal, and temporal lobes of the left hemisphere are shown in (a), (b), (c), and (d), respectively.

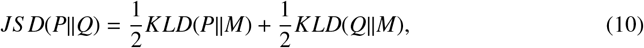

where *JS D*(*P*∥*Q*) is the JSD of distributions *P* and *Q*, the average distribution 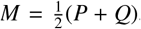, and *KLD*(·∥·) indicates the Kullback–Leibler divergence. The JSD is a symmetric measure for comparing two distributions. A lower JSD value indicates a higher resemblance between the two distributions. To quantify the harmonization performance from the perspective of feature distributions, we computed the JSD of FA and MD distributions across scanners for all four major cortical lobes. The results were summarized in Table 3 and 4 for FA and MD features. We note that our method achieves lower JSD values for both FA and MD features in nearly all cortical regions as compared to results from the LinearRISH method. This reaffirms the efficacy of our method in improving the performance of dMRI harmonization.

**Table 3:**
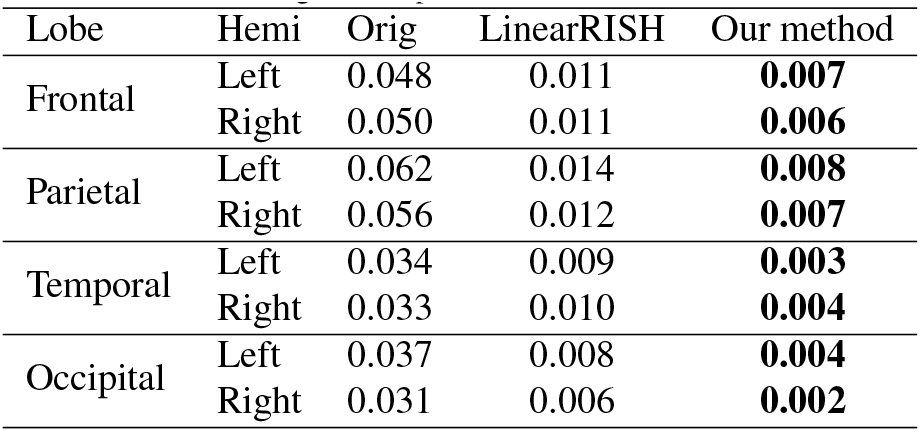
Jensen-Shannon divergence of FA features across scanners before (Orig) and after harmonization (LinearRISH and our method). Results from both left and right hemispheres are listed.

**Table 4:**
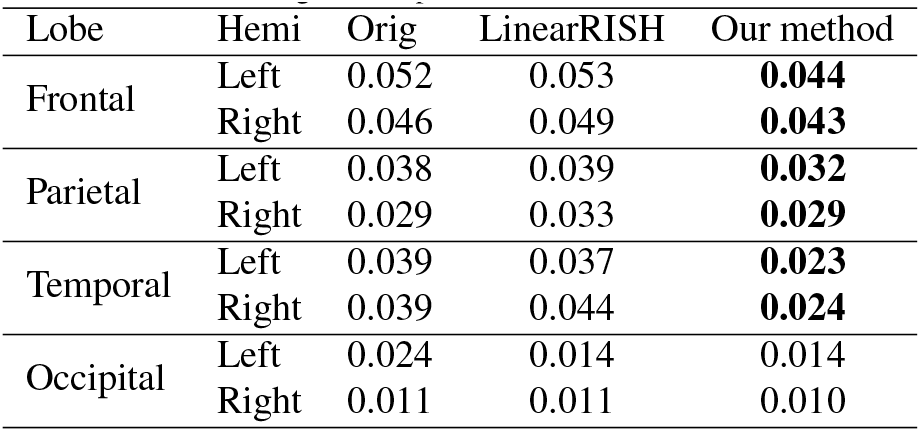
Jensen-Shannon divergence of MD features across scanners before (Orig) and after harmonization (LinearRISH and our method). Results from both left and right hemispheres are listed.

| Lobe | Hemi | Orig | LinearRISH | Our method |
| --- | --- | --- | --- | --- |
| Frontal | Left | 0.052 | 0.053 | **0.044** |
|  | Right | 0.046 | 0.049 | **0.043** |
| Parietal | Left | 0.038 | 0.039 | **0.032** |
|  | Right | 0.029 | 0.033 | **0.029** |
| Temporal | Left | 0.039 | 0.037 | **0.023** |
|  | Right | 0.039 | 0.044 | **0.024** |
| Occipital | Left | 0.024 | 0.014 | 0.014 |
|  | Right | 0.011 | 0.011 | 0.010 |

### 3.7. Preservation of fiber orientation distribution (FOD)

To understand the impact of harmonization on the analysis of structural connectivity, we examined the changes in fiber orientation distribution (FOD) before and after harmonization. First, we estimated the FOD from the original and harmonized dMRI data(Tran & Shi 2015) and extracted the largest FOD peak at each voxel for quantitative comparison. To measure the impact of harmonization on FODs, we computed the cosine similarity of corresponding FOD peak directions estimated from dMRI signals at each voxel before and after harmonization. The average voxel-wise cosine similarity across the white matter was calculated for each subject. For the GE and SIEMENS cohort, the mean and standard deviation of the cosine similarity measure from different harmonization experiments are listed in Table 5. As can be observed from the second and fourth row in Table 5, the proposed method induces minimal changes in the FOD peaks for both cohorts. These results also closely align with those obtained by LinearRISH, shown in the first and third row, which suggests that both our method and LinearRISH are able to preserve the fiber orientations of the original dMRI.

**Table 5:** FOD peaks before and after harmonization. Row 1 and 2: harmonized GE toward SIEMENS; row 3 and 4: harmonized SIEMENS toward GE.

| Dataset | Cosine Similarity |
| --- | --- |
| GE (LinearRISH) | 0.995 ± 0.001 |
| GE (Our Method) | 0.996 ± 0.001 |
| SIEMENS (LinearRISH) | 0.996 ± 0.001 |
| SIEMENS (Our Method) | 0.996 ± 0.001 |

### 3.8. Preservation of sex effects on DTI features

Preserving the inter-subject biological variability in original data is essential during harmonization of data across scanners. To evaluate the impact of harmonization procedures on the preservation of biological variability, we examined sex differences in white matter microstructure properties as such group differences have been found significant and replicable in the baseline cohort of the ABCD study (Lawrence et al. 2023). More specifically, we calculated the effect sizes of sex differences for the GE cohort before and after harmonization toward the SIEMENS cohort. The effect sizes are quantified using Cohen’s d (Cohen 2013):

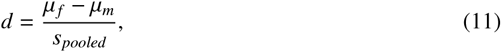

where *μ* _*f*_ and *μ*_*m*_ are the mean feature values for the female and male subjects, respectively. 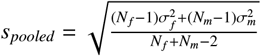 is the pooled standard deviation, where *N* and denote the number of subjects and the standard deviation of a specific gender group.

Considering the sex differences are region-specific (Hsu et al. 2008, Gong et al. 2009, Loópez-Vicente et al. 2021), we examined the effect sizes for all major lobes. The ROIs were delineated by merging the subcortical white matter parcellations extracted by FreeSurfer (Fischl 2012). The mean values of FA in each ROI were computed to reflect the regional white matter maturation. Figure 10 displays the effect sizes of sex differences in FA across brain regions for the GE cohort before and after they were harmonized to the SIEMENS cohort. Compared with LinearRISH, it is evident that our method is more effective in preserving the sex differences after harmonization in most brain regions. It is also notable that the proposed approach is able to preserve effect sizes in all regions with absolute differences (Δ*d*) of effect sizes before and after harmonization no greater than 0.035.

**Figure 10:**
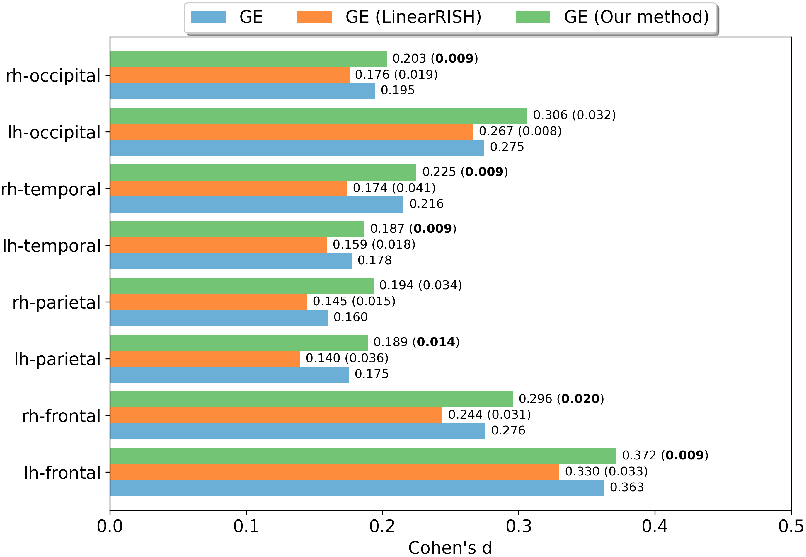
Preservation of sex differences during harmonization. The effect sizes of sex differences in FA before (blue), after harmonization using LinearRISH (orange), and our method (green) for the GE cohort. The quantitative values of effect sizes (*d*) and the absolute differences (Δ*d*) between the effect sizes before and after harmonization are annotated in the form of “*d*(Δ*d*)”.

## 4. Discussions and Conclusions

In this paper, we introduced the personalized idea to dMRI harmonization and comprehensively examined the importance of establishing reliable anatomical correspondences for dMRI harmonization. To mitigate the confounding effects due to the entanglement of inter-subject anatomical variability and site-specific variations in dMRI signals, we presented a personalized template estimation framework and integrated it with RISH features to realize the personalized harmonization of dMRI data. By applying the proposed framework to harmonize dMRI data acquired from Siemens Prisma and GE MR750 scanners in the ABCD study, we demonstrated that our method achieved much improved performance in reducing scanner differences in dMRI signals and preserving sex-related biological variability in original cohorts.

Although the personalized harmonization framework presented in this work employs RISH features for inter-site mapping estimation, the proposed personalized template estimation method is general and can be integrated with other dMRI harmonization techniques for the refinement of anatomical correspondences in different harmonization scenarios. For instance, when there are large variations in the number of gradient directions between the source and target site, the moment-based method (Huynh et al. 2019) might be more preferable as it is independent of the matching of spherical harmonic coefficients. Investigating the effectiveness of embedding the proposed approach in other dMRI harmonization frameworks such as the moment-based method is a direction of our future work.

Besides the conventional frameworks (Mirzaalian et al. 2016, Fortin et al. 2017, Huynh et al. 2019) that explicitly estimate the harmonization mapping function, many learning based methods were developed recently. With scans of traveling subjects obtained from multiple sites, the dMRI harmonization problem could be formulated as a paired image-to-image translation problem (Koppers et al. 2019, Cetin Karayumak et al. 2018, Nath et al. 2019, Hansen et al. 2022). In addition, several unsupervised harmonization methods that do not require explicitly paired scans were proposed for dMRI harmonization (St-Jean et al. 2020, Moyer et al. 2020). While in principle, sophisticated inter-site transformation, learned with deep neural architectures, is beneficial for dMRI harmonization, the inter-subject variability and misalignment of brain anatomy still have significant impact on these methods as the heterogeneity in anatomy (e.g. in cortical gray matter regions) can complicate the learning of inter-site transformation and potentially result in mismatch of training and testing data, which can hinder the reliability of harmonization (Ning et al. 2020, Tax et al. 2019). For our future work, we are also interested in extending our method by developing learning based harmonization algorithms that take into account the inter-subject variability of neuroanatomy.

In the current framework, multiple healthy controls, carefully matched for confounding effects, e.g. age and gender, from each scanning platform are required to avoid bias from confounding variables in the reliable estimation of harmonization mapping functions. For the harmonization of dMRI data from different cohorts with a limited overlap of related confounding variables such as age, our method can be extended to consider these confounding variables as additive terms in a regression model. Similar to the ComBat method (Fortin et al. 2016), the reference features can be adjusted by regressing out the confounding variables to reveal site-specific characteristics for harmonization. This possible extension will be another direction of our future work.

## Acknowledgement

This work was supported by National Institute of Health (NIH) under grants R01EB022744, RF1AG056573, RF1AG064584, R21AG064776, P41EB015922, U19AG078109.

## Data and code availability

The scripts and code used in this study will be shared publicly upon publication of the article. The data used in this paper are part of the Adolescent Brain Cognitive Development (ABCD) Data Release 4.0 from the NIMH Data Archive (NDA).

## References

Andersson, J. L., Skare, S., & Ashburner, J., 2003. How to correct susceptibility distortions in spin-echo echo-planar images: application to diffusion tensor imaging. NeuroImage, 20 (2), 870–888.

Andersson, J. L., Xu, J., Yacoub, E., Auerbach, E., Moeller, S., & Ugurbil, K., 2012. A comprehensive gaussian process framework for correcting distortions and movements in diffusion images. In in Proc. Annu. Meeting Int. Soc. Magn. Reson. Med.

Avants, B. B., Tustison, N., & Song, G., 2009. Advanced normalization tools (ants). Insight J.

Basser, P. J., Mattiello, J., & LeBihan, D., 1994. MR diffusion tensor spectroscopy and imaging. Biophys. J., 66 (1), 259–267.

Casey, B. J., Cannonier, T., Conley, M. I., Cohen, A. O., Barch, D. M., Heitzeg, M. M., Soules, M. E., Teslovich, T., Dellarco, D. V., Garavan, H., Orr, C. A., Wager, T. D., Banich, M. T., Speer, N. K., Sutherland, M. T., Riedel, M. C., Dick, A. S., Bjork, J. M., Thomas, K. M., Chaarani, B., Mejia, M. H., Hagler, D. J., Daniela Cornejo, M., Sicat, C. S., Harms, M. P., Dosenbach, N. U., Rosenberg, M., Earl, E., Bartsch, H., Watts, R., Polimeni, J. R., Kuperman, J. M., Fair, D. A., & Dale, A. M., 2018. The adolescent brain cognitive development (abcd) study: Imaging acquisition across 21 sites. Dev. Cogn. Neurosci., 32, 43–54.

Cetin Karayumak, S., Bouix, S., Ning, L., James, A., Crow, T., Shenton, M., Kubicki, M., & Rathi, Y., 2019. Retrospective harmonization of multi-site diffusion MRI data acquired with different acquisition parameters. NeuroImage, 184, 180–200.

Cetin Karayumak, S., Kubicki, M., & Rathi, Y., 2018. Harmonizing diffusion mri data across magnetic field strengths. In in Proc. Int. Conf. Med. Image Comput. Comput. Interv. 116–124. Springer.

Cohen, J., 2013. Statistical power analysis for the behavioral sciences. Academic press.

Diamond, S. & Boyd, S., 2016. Cvxpy: A python-embedded modeling language for convex optimization. J. Mach. Learn. Res., 17 (1), 2909–2913.

Domahidi, A., Chu, E., & Boyd, S., 2013. Ecos: An socp solver for embedded systems. In 2013 European control conference (ECC) 3071–3076. IEEE.

Fischl, B., 2012. Freesurfer. NeuroImage, 62 (2), 774–781.

Fortin, J. P., Parker, D., Tunç, B., Watanabe, T., Elliott, M. A., Ruparel, K., Roalf, D. R., Satterthwaite, T. D., Gur, R. C., Gur, R. E., Schultz, R. T., Verma, R., & Shinohara, R. T., 2017. Harmonization of multi-site diffusion tensor imaging data. NeuroImage, 161, 149–170.

Fortin, J. P., Sweeney, E. M., Muschelli, J., Crainiceanu, C. M., & Shinohara, R. T., 2016. Removing inter-subject technical variability in magnetic resonance imaging studies. NeuroImage, 132, 198–212.

Gong, G., Rosa-Neto, P., Carbonell, F., Chen, Z. J., He, Y., & Evans, A. C., 2009. Age-and gender-related differences in the cortical anatomical network. Journal of Neuroscience, 29 (50), 15684–15693.

Hagler Jr, D. J., Ahmadi, M. E., Kuperman, J., Holland, D., McDonald, C. R., Halgren, E., & Dale, A. M., 2009. Automated white-matter tractography using a probabilistic diffusion tensor atlas: Application to temporal lobe epilepsy. Human brain mapping, 30 (5), 1535–1547.

Hagler Jr, D. J., Hatton, S., Cornejo, M. D., Makowski, C., Fair, D. A., Dick, A. S., Sutherland, M. T., Casey, B. J., Barch, D. M., Harms, M. P., Watts, R., Bjork, J. M., Garavan, H. P., Hilmer, L., Pung, C. J., Sicat, C. S., Kuperman, J., Bartsch, H., Xue, F., Heitzeg, M. M., et al., 2019. Image processing and analysis methods for the adolescent brain cognitive development study. Neuroimage, 202, 116091.

Hansen, C. B., Schilling, K. G., Rheault, F., Resnick, S., Shafer, A. T., Beason-Held, L. L., & Landman, B. A., 2022. Contrastive semi-supervised harmonization of single-shell to multi-shell diffusion mri. Magn. Reson. Imaging, 93, 73–86.

Hsu, J.-L., Leemans, A., Bai, C.-H., Lee, C.-H., Tsai, Y.-F., Chiu, H.-C., & Chen, W.-H., 2008. Gender differences and age-related white matter changes of the human brain: a diffusion tensor imaging study. Neuroimage, 39 (2), 566–577.

Huynh, K. M., Chen, G., Wu, Y., Shen, D., & Yap, P. T., 2019. Multi-Site Harmonization of Diffusion MRI Data via Method of Moments. IEEE Trans. Med. Imaging, 38 (7), 1599–1609.

Johnson, W. E., Li, C., & Rabinovic, A., 2007. Adjusting batch effects in microarray expression data using empirical Bayes methods. Biostatistics, 8 (1), 118–127.

Kawin Setsompop 1, Gagoski, B. A., Polimeni, J. R., Witzel, T., Wedeen, V. J., & Wald, L. L., 2011. Blipped-controlled aliasing in parallel imaging for simultaneous multislice echo planar imaging with reduced g-factor penalty. Magnetic Resonance in Medicine, 67 (5), 1210–1224.

Koppers, S., Bloy, L., Berman, J. I., Tax, C. M., Edgar, J. C., & Merhof, D., 2019. Spherical Harmonic Residual Network for Diffusion Signal Harmonization. In in Proc. Int. Conf. Med. Image Comput. Comput. Interv. 173–182.

Lawrence, K. E., Abaryan, Z., Laltoo, E., Hernandez, L. M., Gandal, M. J., McCracken, J. T., & Thompson, P. M., 2023. White matter microstructure shows sex differences in late childhood: Evidence from 6797 children. Human Brain Mapping, 44 (2), 535–548.

Leek, J. T. & Storey, J. D., 2007. Capturing heterogeneity in gene expression studies by surrogate variable analysis. PLoS Genet., 3 (9), 1724–1735.

Lin, J., 1991. Divergence measures based on the shannon entropy. IEEE Transactions on Information theory, 37 (1), 145–151.

Loópez-Vicente, M., Lamballais, S., Louwen, S., Hillegers, M., Tiemeier, H., Muetzel, R. L., & White, T., 2021. White matter microstructure correlates of age, sex, handedness and motor ability in a population-based sample of 3031 school-age children. Neuroimage, 227, 117643.

Magnotta, V. A., Matsui, J. T., Liu, D., Johnson, H. J., Long, J. D., Bolster, B. D., Mueller, B. A., Lim, K., Mori, S., Helmer, K. G., Turner, J. A., Reading, S., Lowe, M. J., Aylward, E., Flashman, L. A., Bonett, G., & Paulsen, J. S., 2012. MultiCenter reliability of diffusion tensor imaging. Brain Connect., 2 (6), 345–355.

Manjón, J. V., Coupé, P., Concha, L., Buades, A., Collins, D. L., & Robles, M., 2013. Diffusion weighted image denoising using overcomplete local pca. PloS One, 8 (9), 1–12.

Mirzaalian, H., Ning, L., Savadjiev, P., Pasternak, O., Bouix, S., Michailovich, O., Karmacharya, S., Grant, G., Marx, C. E., Morey, R. A., Flashman, L. A., George, M. S., McAllister, T. W., Andaluz, N., Shutter, L., Coimbra, R., Zafonte, R. D., Coleman, M. J., Kubicki, M., Westin, C. F., Stein, M. B., Shenton, M. E., & Rathi, Y., 2018. Multisite harmonization of diffusion MRI data in a registration framework. Brain Imaging Behav., 12 (1), 284–295.

Mirzaalian, H., Ning, L., Savadjiev, P., Pasternak, O., Bouix, S., Michailovich, O., Marx, C. E., Morey, R. A., Flashman, L. A., George, M. S., Mcallister, T. W., Shutter, L., Coimbra, R., Zafonte, R. D., Coleman, M. J., Kubicki, M., Westin, C. F., Stein, M. B., Shenton, M. E., & Rathi, Y., 2016. Inter-site and inter-scanner diffusion MRI data harmonization. NeuroImage, 135, 311–323.

Mirzaalian, H., Pierrefeu, A. D., Savadjiev, P., Pasternak, O., Bouix, S., & Kubicki, M., 2015. Harmonizing diffusion mri data across multiple sites and scanners. In in Proc. Int. Conf. Med. Image Comput. Comput. Interv. 12–19.

Moyer, D., Ver Steeg, G., Tax, C. M., & Thompson, P. M., 2020. Scanner invariant representations for diffusion mri harmonization. Magn. Reson. Med., 84 (4), 2174–2189.

Mueller, S. G., Weiner, M. W., Thal, L. J., Petersen, R. C., Jack, C., Jagust, W., Trojanowski, J. Q., Toga, A. W., & Beckett, L., 2005. The alzheimer’s disease neuroimaging initiative. Neuroimaging Clin. N. Am., 15 (4), 869–877.

Nath, V., Remedios, S., Parvathaneni, P., Hansen, C., Bayrak, R., Bermudez, C., Blaber, J., Schilling, K., Janve, V., Gao, Y., & Huo, Y., 2019. Harmonizing 1.5t/3t diffusion weighted mri through development of deep learning stabilized microarchitecture estimators. In in Proc. SPIE. Medical Imaging 173–182.

Ning, L., Bonet-Carne, E., Grussu, F., Sepehrband, F., Kaden, E., Veraart, J., Blumberg, S. B., Khoo, C. S., Palombo, M., Kokkinos, I., Alexander, D. C., Coll-Font, J., Scherrer, B., Warfield, S. K., Karayumak, S. C., Rathi, Y., Koppers, S., Weninger, L., Ebert, J., Merhof, D., Moyer, D., Pietsch, M., Christiaens, D., Gomes Teixeira, R. A., Tournier, J.-D., Schilling, K. G., Huo, Y., Nath, V., Hansen, C., Blaber, J., Landman, B. A., Zhylka, A., Pluim, J. P. W., Parker, G., Rudrapatna, U., Evans, J., Charron, C., Jones, D. K., & Tax, C. M. W., 2020. Cross-scanner and cross-protocol multi-shell diffusion MRI data harmonization: Algorithms and results. NeuroImage, 221, 117–128.

St-Jean, S., Viergever, M. A., & Leemans, A., 2020. Harmonization of diffusion mri data sets with adaptive dictionary learning. Hum. Brain Mapp., 41 (16), 4478–4499.

Tax, C. M. W., Grussu, F., Kaden, E., Ning, L., Rudrapatna, U., Evans, C., St-Jean, S., Leemans, A., Koppers, S., Merhof, D., & Ghosh, A., 2019. Cross-scanner and cross-protocol diffusion mri data harmonisation: A benchmark database and evaluation of algorithms. NeuroImage, 195, 285–299.

Thompson, P. M., Schwartz, C., Lin, R. T., Khan, A. A., & Toga, A. W., 1996. Three-dimensional statistical analysis of sulcal variability in the human brain. J. Neurosci., 16 (13), 4261–4274.

Tran, G. & Shi, Y., 2015. Fiber orientation and compartment parameter estimation from multi-shell diffusion imaging. IEEE Trans. Med. Imaging, 34 (11), 2320–2332.

Uylings, H. B., Rajkowska, G., Sanz-Arigita, E., Amunts, K., & Zilles, K., 2005. Consequences of large interindividual variability for human brain atlases: Converging macroscopical imaging and microscopical neuroanatomy. Anat. Embryol., 210 (5-6), 423–431.

Van Essen, D. C., Smith, S. M., Barch, D. M., Behrens, T. E., Yacoub, E., Ugurbil, K., & Consortium, W.-M. H., 2013. The wu-minn human connectome project: an overview. NeuroImage, 80, 62–79.

Vollmar, C., O’Muircheartaigh, J., Barker, G. J., Symms, M. R., Thompson, P., Kumari, V., Duncan, J. S., Richardson, M. P., & Koepp, M. J., 2010. Identical, but not the same: Intra-site and inter-site reproducibility of fractional anisotropy measures on two 3.0t scanners. NeuroImage, 51 (4), 1384–1394.

Xue, W., Mou, X., Zhang, L., Bovik, A. C., & Feng, X., 2014. Blind image quality assessment using joint statistics of gradient magnitude and laplacian features. IEEE Trans. Image Process., 23 (11), 4850–4862.

Zhu, T., Hu, R., Qiu, X., Taylor, M., Tso, Y., Yiannoutsos, C., Navia, B., Mori, S., Ekholm, S., Schifitto, G., & Zhong, J., 2011. Quantification of accuracy and precision of multi-center dti measurements: A diffusion phantom and human brain study. NeuroImage, 56 (3), 1398–1411.

Zhuang, J., Hrabe, J., Kangarlu, A., Xu, D., Bansal, R., Branch, C. A., & Peterson, B. S., 2006. Correction of eddy-current distortions in diffusion tensor images using the known directions and strengths of diffusion gradients. Journal of Magnetic Resonance Imaging: An Official Journal of the International Society for Magnetic Resonance in Medicine, 24 (5), 1188–1193.

